# Regulatory T-cells are required for neonatal heart regeneration

**DOI:** 10.1101/355065

**Authors:** Jiatao Li, Kevin Y. Yang, Rachel Chun Yee Tam, Vicken W. Chan, Bao Sheng, Shohei Hori, Bin Zhou, Yuk Ming Dennis Lo, Kathy O. Lui

**Affiliations:** Department of Chemical Pathology, Prince of Wales Hospital, The Chinese University of Hong Kong, Hong Kong, China; Li Ka Shing Institute of Health Sciences, Prince of Wales Hospital, The Chinese University of Hong Kong, Hong Kong, China; Centre for Integrative Medical Sciences, RIKEN, Japan; The State Key Laboratory of Cell Biology, CAS Center for Excellence in Molecular Cell Science, Shanghai Institute of Biochemistry and Cell Biology, Chinese Academy of Sciences, University of Chinese Academy of Sciences, Shanghai, China

**Keywords:** CD4^+^ regulatory T-cells, heart regeneration, cardiomyocytes, cardiac fibrosis, macrophages, single-cell transcriptomics

## Abstract

Previous work has elegantly demonstrated that, unlike adult mammalian heart, the neonatal heart is able to regenerate after injury from postnatal day (P) 1 to 7. Recently, macrophages are found to be required in the repair process as depletion of which abolishes endogenous regenerative capability of the neonatal heart. Nevertheless, whether innate immunity alone is sufficient for neonatal heart regeneration is obscure. Here, we investigate a hitherto novel role of FOXP3^+^ regulatory T-cells (Treg) in neonatal heart regeneration. Unlike their wild type counterparts, NOD/SCID mice that are deficient for T-cells but innate immune cells including macrophages fail to regenerate their injured heart as early as P3. In wild type mice, both conventional CD4^+^ T-cells and Treg are recruited to cardiac muscle within the first week after injury. Treatment with the lytic anti-CD4 antibody that specifically depletes conventional CD4^+^ T-cells leads to reduced cardiac fibrosis; while treatment with the lytic anti-CD25 antibody that specifically depletes CD4^+^CD25^hi^FOXP3^+^ Treg contributes to increased fibrosis of the neonatal heart after injury. Moreover, adoptive transfer of Treg to NOD/SCID mice results in mitigated fibrosis and increased proliferation and function of cardiac muscle of the neonatal heart after injury. Mechanistically, single cell transcriptomic profiling reveals that Treg are a source of chemokines and cytokines that attract monocytes and macrophages previously known to drive neonatal heart regeneration. Furthermore, Treg directly promote proliferation of both mouse and human cardiomyocytes in a paracrine manner. Our findings uncover an unappreciated mechanism in neonatal heart regeneration; and offer new avenues for developing novel therapeutics targeting Treg-mediated heart regeneration.

## Introduction

Heart disease remains the leading cause of deaths worldwide. Yet the mammalian heart is notorious for its inability to repair and regenerate after injury; and the loss of cardiac muscle is replaced by scar tissues that further compromise heart function. Intriguingly, recent studies demonstrate that the mouse heart can transiently regenerate after birth till postnatal day 7 (P7) in a range of injury models including amputation of the ventricular apex via apical resection (AR)^1^, myocardial infarction (MI)^2^, cryoinfarction (CI)^3^ and cardiomyocyte-specific cell death^4^. Importantly, neonatal heart regeneration is also observed in human^5^. In contrast to adults, neonatal cardiomyocytes can proliferate so the heart muscle is regenerated after injury with robust angiogenesis and minimal fibrosis^1,6^. Nevertheless, mechanisms driving neonatal heart regeneration are largely elusive. Since decades ago, the immune system has been known to orchestrate tissue repair as immune cells regulate both angiogenesis and fibrosis. Therefore, understanding how immune cells participate in neonatal heart regeneration would shed light on development of potential therapeutics to promote heart repair and regeneration.

During the last decade, innate immunity, particularly macrophages and their various polarization states, have been considered as a central regulator of the tissue healing processes. Indeed, previous studies demonstrate that macrophages are required in neonatal heart regeneration^4,7^. Nonetheless, the role of adaptive immunity in neonatal heart regeneration has not been investigated. Long recognized as potent suppressors of the immune system, CD4^+^ regulatory T-cells (Treg) are recently rediscovered as direct or indirect regulators of organ regeneration^8^. We and others have showed that Treg promote repair of skeletal muscle^9,10^, skin^11^, lung^12^, bone^13^, central nervous system^14^ and peripheral vascular system^15^ after injury. Treg are recruited in response to neoantigens to damaged tissues for resolution of inflammation and regulation of innate immune response^16^. Mechanistically, Treg can directly activate skeletal muscle progenitors/satellite cells via amphiregulin^9^ or stimulate hair follicle stem cells via Notch signaling^11^. Moreover, we have also reported that Treg directly promote endothelial cell proliferation and indirectly inhibit activation of conventional CD4^+^ T-cells that impair vascular regeneration after ischemic injury in type-2 diabetes^15^.

In this study, we further investigate the unappreciated role of Treg in neonatal heart regeneration. We utilized CI as a major injury model because the fibrotic responses during neonatal heart repair resemble the healing processes of the adult heart. Unlike their wild type counterparts, T-cell deficient NOD/SCID mice failed to regenerate their injured myocardium at P3. Treg were recruited to cardiac muscle within the first week after injury. Treatment with the anti-CD25 antibody (clone PC61) that specifically depletes Treg led to increased cardiac fibrosis; while treatment with the anti-CD4 antibody (clone GK1.5) that specifically depletes conventional CD4^+^ T-cells reduced fibrosis in wild type neonatal hearts after injury. Our results of adoptive transfer experiments also indicate that Treg promoted neonatal heart regeneration in NOD/SCID mice after injury. Single cell transcriptomic profiling reveals that Treg are chemoattractive to innate immune cells including monocytes or macrophages. Furthermore, Treg directly promote human and mouse cardiomyocyte proliferation in a paracrine manner.

## Results

### Innate immunity alone is insufficient for neonatal heart regeneration of NOD/SCID mice

Previous study shows that depletion of macrophages leads to excessive fibrosis and lack of neoangiogenesis after injury, resulting in compromised neonatal heart regeneration^7^. Nevertheless, whether innate immune cells including macrophages are sufficient for neonatal heart regeneration is unanswered. To address this question, we induced CI as previously described^3,17^ to P3 (regenerating) or P8 (non-regenerating) neonatal mouse heart of NOD/SCID that harbors innate immune cells such as macrophages without functional adaptive immune cells such as T-cells. We then examined heart regeneration at day 28 post injury (Fig 1A). We used NOD/ShiLtJ (NOD) as a control of NOD/SCID because the latter is its descendant; and autoimmune diabetes does not occur during the study period (i.e. before 4-8 weeks of age). We also included ICR as an additional control because NOD is its descendant but ICR does not develop autoimmunity. Our results showed that the P3 heart of both NOD and ICR regenerated better than their P8 counterparts (Fig 1B), consistent with previous reports. However, both the P3 and P8 hearts of NOD/SCID did not regenerate (Fig 1B). We then quantified the degree of injury by Masson’s trichrome staining that identifies collagen fibers. The P3 heart of NOD (Fig 1C, D) and ICR (Fig 1E, F) had significantly less fibrotic tissues than their P8 hearts, respectively. Compared to that of NOD, however, the P3 heart of NOD/SCID showed significantly more excessive scar tissue formation (Fig 1G, H), suggestive of impaired neonatal heart repair. Furthermore, we also examined heart function at 2 months post injury by echocardiography (Fig 1I). Our data revealed that the P3 heart of NOD fully regenerated functionally as there was no significant difference in %fractional shortening between the CI and sham groups (Fig 1J). On the other hand, the P3 heart of NOD/SCID showed significantly less functional regeneration by comparing with its sham control or the P3 heart of NOD (Fig 1J). Taken together, our findings showed that innate immune cells alone in NOD/SCID mice were insufficient for driving functional neonatal heart regeneration.

**Fig. 1.**
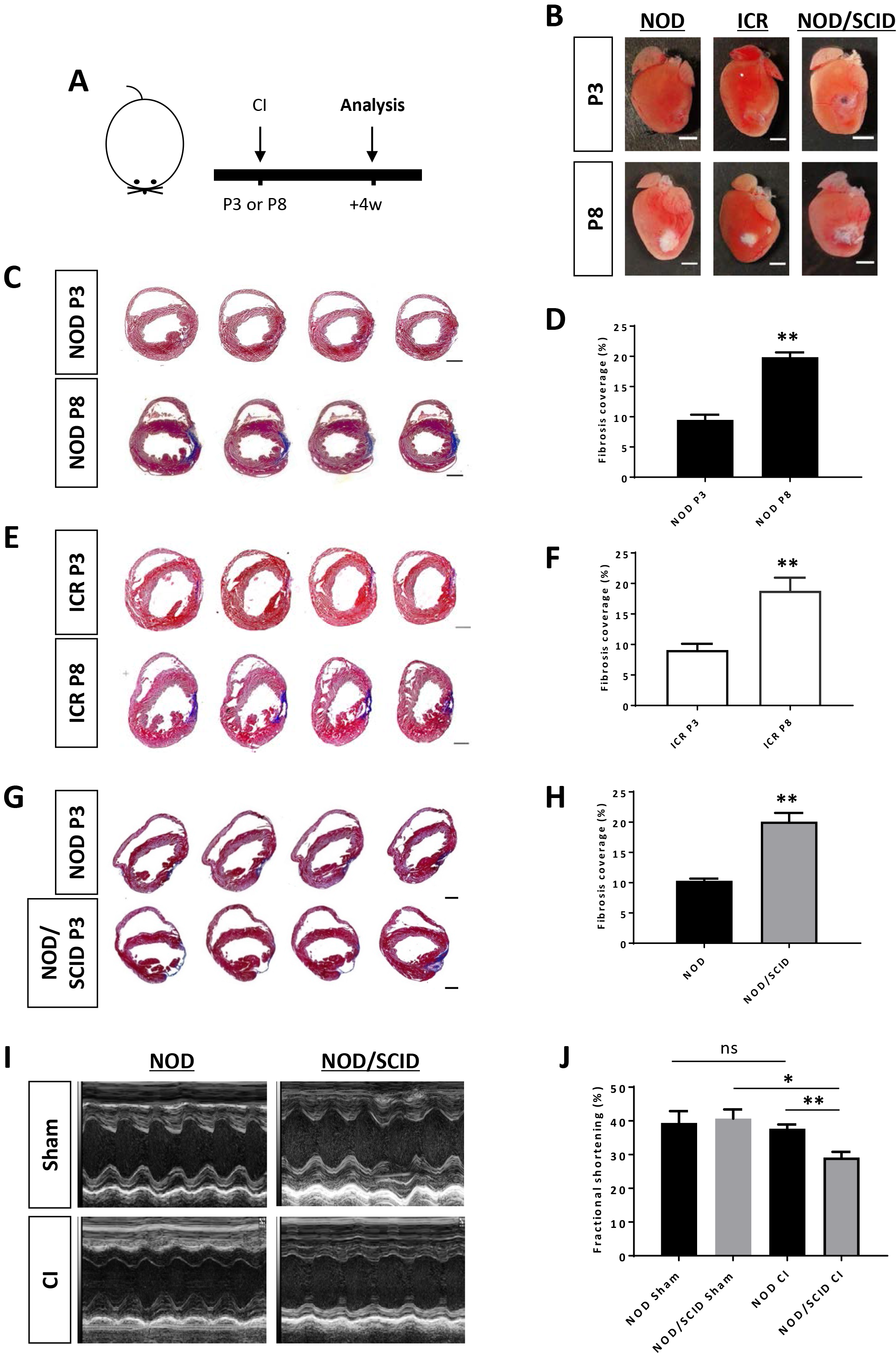
The neonatal heart of NOD/SCID fails to regenerate. (A) Schematic diagram showing experimental design. (B) Images of scar tissues at 4 weeks post CI to P3 or P8 hearts, scale bars: 2000 um. (C, E, G) Masson’s trichrome staining showing representative serial cross sections of fibrotic tissues in blue at 4 weeks post CI, scale bars: 1000 um. (D, F, H) Quantification of fibrotic tissue coverage based on (C, E, G respectively). (I) Echocardiographic analysis and (J) quantification showing %fractional shortening at 8 weeks post CI. (D, F, H, J) Data are presented as mean±S.E.M., *P<0.05, **P<0.01, n=6 per group.

### Loss-of-function of conventional CD4^+^ T-cells leads to reduced cardiac fibrosis

To ask if T-cells are recruited to the neonatal heart after injury, we first performed immunostaining for CD3 in the border zone at day 1 following CI to a P3 ICR heart. Our results demonstrated infiltration of CD3^+^ T-cells (Fig 2A). We then asked which subsets of CD3^+^ T-cells responsible for neonatal heart regeneration as the absence of which in NOD/SCID mice contributed to impaired regeneration. We quantified infiltration of CD4^+^ or CD8^+^ T-cells after CI by flow cytometry. We found that there was a significant increase in absolute number of CD3^+^CD4^+^ T-cells at days 7 and 14 after CI to a P8 heart compared to its sham control; yet there was no significant difference in absolute number of CD3^+^CD4^+^ T-cells within the first two weeks after CI to a P3 heart compared to sham (Fig 2B). Moreover, we did not observe significant difference in absolute number of CD3^+^CD8^+^ T-cells within the first two weeks after CI to a P3 or P8 heart compared to sham (Fig 2C). Our findings indicated a potential role of CD4^+^ T-cells in determining the outcome of heart regeneration as they significantly infiltrated into the non-regenerating P8 heart but not the regenerating P3 heart after CI. To test this hypothesis, we respectively depleted CD4^+^ and CD8^+^ T-cells using the lytic anti-CD4 (clone GK1.5) and anti-CD8 (clone YTS169) antibodies after CI to a P8 heart and examined heart regeneration at day 28 post CI (Fig 2D). We confirmed highly efficient depletion by flow cytometry as CD3^+^CD4^+^ (Fig S1A) and CD3^+^CD8^+^ (Fig S1B) T-cells were rarely present in the peripheral blood at day 28 after the use of respective antibody. Intriguingly, Masson’s trichrome staining demonstrated significantly reduced cardiac fibrosis in the GK1.5-treated compared to the untreated control or YTS169-treated group (Fig 2E, F). Moreover, results of immunostaining showed that scar tissue of the control or YTS169-treated group was rich in type-1 collagen (COLA1, Fig 2G). To ask if GK1.5 improved heart regeneration by facilitating proliferation of cardiomyocytes, we performed immunostaining for proliferation marker (Ki67) and cardiac troponin T (cTnT) at the border zone. Our results revealed that there was significantly increased number of Ki67^+^cTNT^+^ cells in the GK1.5-treated compared to untreated control group (Fig 2H). Together, depletion of CD4^+^ T-cells promoted regeneration of the P8 heart after injury.

**Fig. 2.**
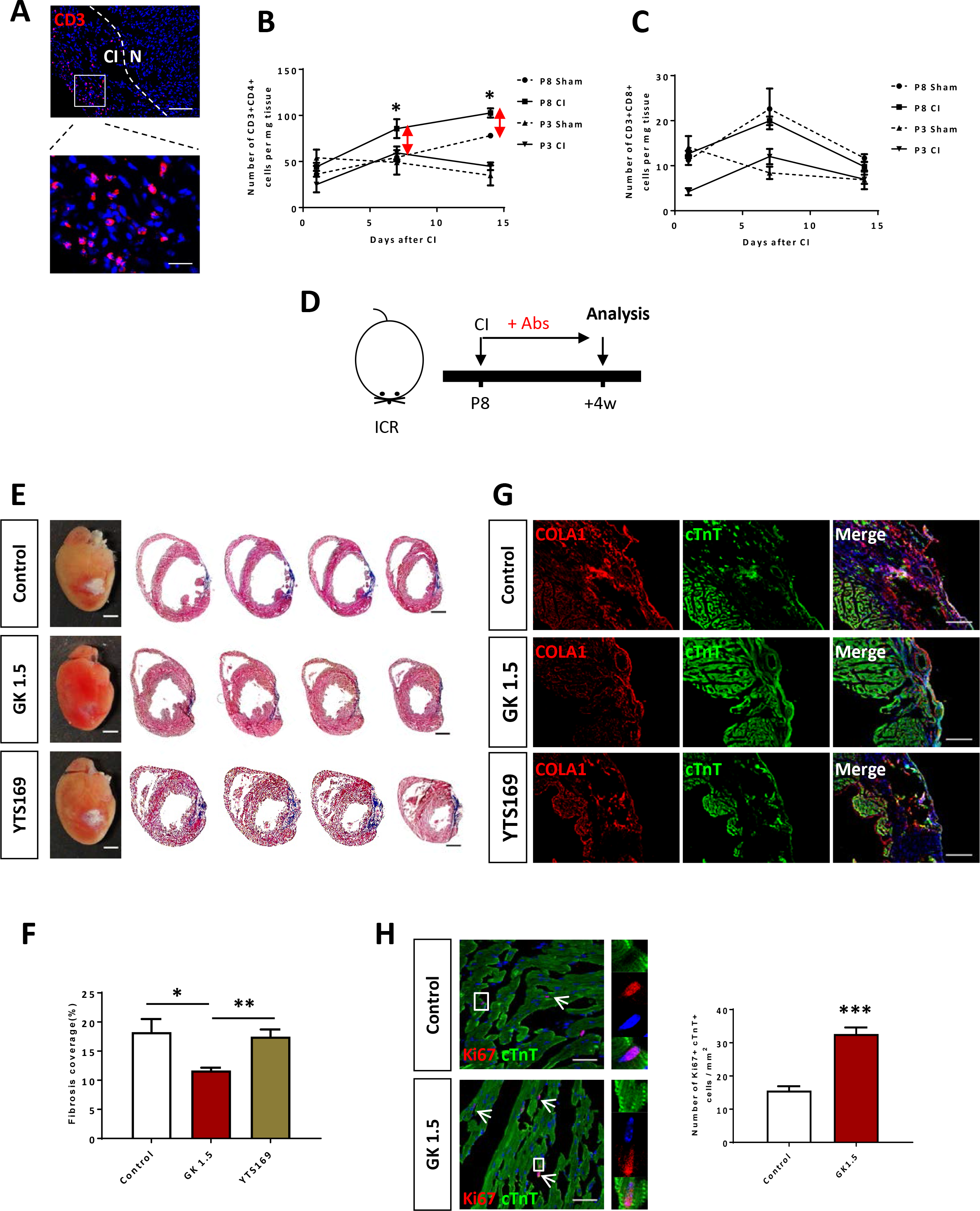
Recruitment and function of CD4^+^ and CD8^+^ T-cells during neonatal heart regeneration. (A) Immunostaining on frozen sections for CD3^+^ (red) cells with nuclear DAPI counterstain (blue) showing infiltration of T-cells into infarcted (CI) and normal (N) myocardium at day 1 post CI to a P3 heart, scale bar: 100 um. Square denotes enlarged image, scale bar: 50 um. Quantification of flow cytometric analyses showing absolute number of (B) CD3^+^CD4^+^ or (C) CD3^+^CD8^+^ T-cells per mg tissue within the first 2 weeks post CI to a P3 or P8 heart compared to sham. (D) Schematic diagram showing experimental design and analysis performed at 4 weeks post CI. (E) Images of scar tissues, scale bars: 2000 um; and Masson’s trichrome staining showing representative serial cross sections of fibrotic tissues in blue, scale bars: 1000 um. (F) Quantification of fibrotic tissue coverage based on (E). Immunostaining on frozen sections for (G) COLA1^+^ (red) and cTnT^+^ (green) cells within the infarct zone or (H) Ki67^+^ (red) and cTnT^+^ (green) cells within the border zone. (H) Quantification of absolute number of Ki67^+^cTnT^+^ cardiomyocytes per mm^2^ area. Arrows indicate cardiomyocytes positive for Ki67 and square denotes magnified images on the right. (B, C, F, H) Data are presented as mean±S.E.M., *P<0.05, **P<0.01, ***P<0.001. (B, C) n=4, (F) n=8 or (H) n=6 per group.

### Loss-of-function of FOXP3^+^ Treg contributes to increased cardiac fibrosis of the neonatal heart

We found that treatment with GK1.5 significantly reduced both the %CD4^+^FOXP3^−^ and %CD4^+^FOXP3^+^ populations; and significantly enhanced the %CD4−FOXP3+ population (Fig S1C). To ask if FOXP3^+^ Treg play a role in neonatal heart regeneration, we backcrossed the *Foxp3*^hCD2^ reporter “knockin” allele^18^ onto NOD background, allowing us to purify FOXP3^+^ Treg via their surface expression of hCD2. We first quantified infiltration of CD3^+^CD4^+^hCD2^+^ Treg into the damaged myocardium after CI to a P3 heart by flow cytometry (Fig 3A). There was a significant increase in %CD4^+^FOXP3^+^ Treg among CD4^+^ T-cells (Fig 3B) or in absolute number of CD4^+^FOXP3^+^ Treg per mg tissue (Fig 3C) at day 7 post CI compared to sham. We then depleted CD4^+^CD25^hi^FOXP3^+^ Treg via the lytic anti-CD25 antibodies (clone PC61) as previously described (Fig 3D, Fig S2)^15,19^. Treatment with PC61 significantly increased cardiac fibrosis (Fig 3E, F) and COLA1 deposition (Fig 3G); and significantly reduced cTnT^+^ myocardium in the infarct zone (Fig 3G, H) and lessened Ki67^+^cTnT^+^ proliferating cardiomyocytes in the border zone at day 28 post CI (Fig 3I, J). Altogether, our loss-of-function experiments showed that ablation of Treg impaired the endogenous regenerative capability of a P3 heart after CI.

**Fig. 3.**
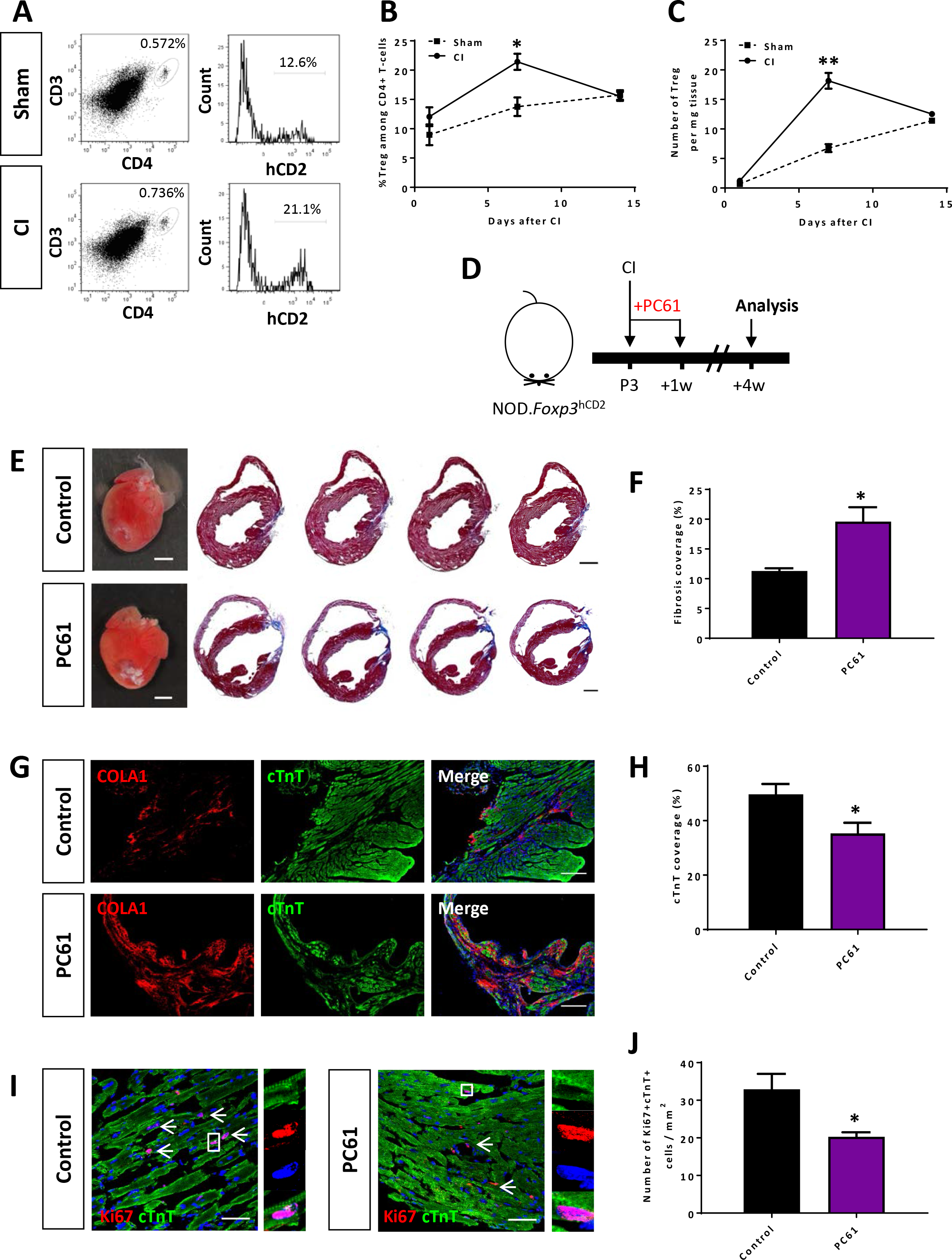
Loss-of-function of CD4^+^ Treg leads to increased fibrosis and reduced cardiomyocyte proliferation after cryoinfarction. (A) Flow cytometric analysis showing infiltration of CD3^+^CD4^+^hCD2^+^ Treg into the myocardium at day 7 post CI to a P3 heart compared to sham. Quantification of (B) %Treg among total CD4^+^ T-cells or (C) absolute number of Treg per mg tissue within the first 2 weeks post CI based on (A). (D) Schematic diagram showing experimental design and analysis performed at 4 weeks post CI. (E) Images of scar tissues, scale bars: 2000 um; and Masson’s trichrome staining showing representative serial cross sections of fibrotic tissues in blue, scale bars: 1000 um. (F) Quantification of fibrotic tissue coverage based on (E). Immunostaining on frozen sections for (G) COLA1^+^ (red) and cTnT^+^ (green) cells within the infarct zone or (I) Ki67^+^ (red) and cTnT^+^ (green) cells within the border zone. Quantification of absolute number of (H) %cTnT^+^ coverage or (J) Ki67^+^cTnT^+^ cardiomyocytes per mm^2^ area. (I) Arrows indicate cardiomyocytes positive for Ki67 and square denotes magnified images on the right. (B, C, F, H, J) Data are presented as mean±S.E.M., *P<0.05, **P<0.01. (B, C) n=4 or (F, H, J) n=8 per group.

### Adoptive transfer of FOXP3^+^ Treg promotes neonatal heart regeneration of NOD/SCID mice

To further confirm the role of Treg in neonatal heart regeneration, we purified 1 million FOXP3^+^ (hCD2^+^) Treg from the spleen of NOD.*Foxp3*^hCD2^ mice for adoptive transfer via intraperitoneal injection as previously described^20^ into NOD/SCID mice so the recipients only harbored Treg without other conventional T-cells (Fig 4A). By flow cytometric analysis on day 7 following adoptive transfer, we found that about 88% of total CD3^+^CD4^+^ T-cells were hCD2^+^ Treg in the spleen and heart, respectively (Fig 4B). Moreover, we also quantified engraftment of CD3^+^CD4^+^hCD2^+^ Treg weekly during the first 4 weeks following adoptive transfer and found that the %FOXP3^+^ Treg among total splenocytes increased gradually and reached about 5% by day 28 post adoptive transfer (Fig 4C), similar to a wild type mouse.

**Fig. 4.**
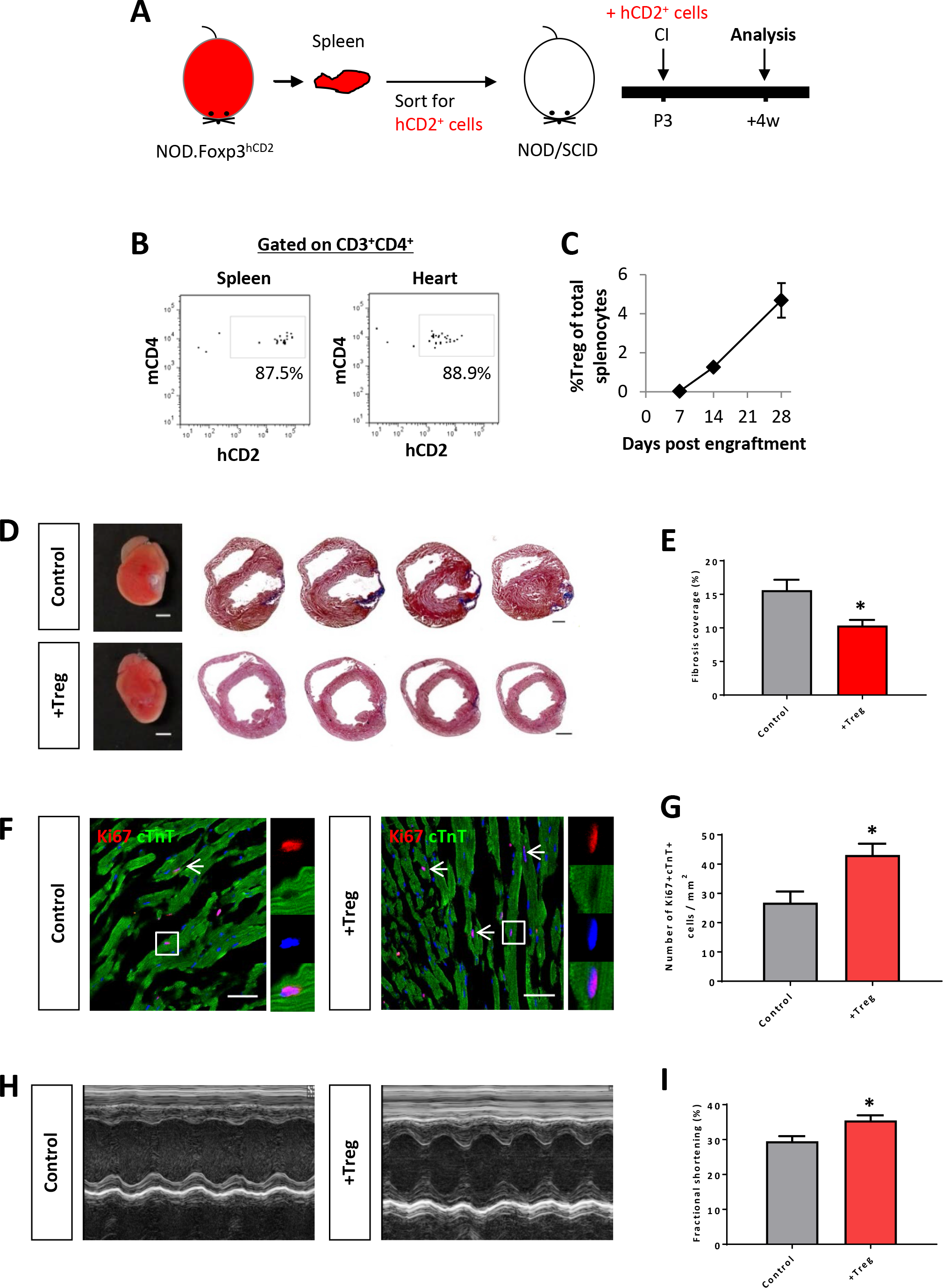
Adoptive transfer of CD4^+^ Treg potentiates neonatal heart regeneration after cryoinfarction. (A) Schematic diagram showing strategy of adoptive transfer. (B) Flow cytometric analysis showing engraftment and infiltration of CD3^+^CD4^+^hCD2^+^ Treg in the spleen and myocardium, respectively, at day 7 following CI to a P3 heart of NOD/SCID mice. (C) Quantification of %Treg among total splenocytes at days 7, 14 and 28 after adoptive transfer by flow cytometry. (D) Images of scar tissues, scale bars: 2000 um; and Masson’s trichrome staining showing representative serial cross sections of fibrotic tissues in blue at 4 weeks post CI, scale bars: 1000 um. (E) Quantification of fibrotic tissue coverage based on (D). (F) Immunostaining on frozen sections for Ki67^+^ (red) and cTnT^+^ (green) cells within the border zone at 4 weeks post CI. Arrows indicate cardiomyocytes positive for Ki67 and square denotes magnified images on the right. (G) Quantification of absolute number of Ki67^+^cTnT^+^ cardiomyocytes per mm^2^ area. (H) Echocardiographic analysis and (I) quantification showing %fractional shortening at 4 weeks post CI. (C, E, G, I) Data are presented as mean±S.E.M., *P<0.05. (C) n=4 or (E, G, I) n=6 per group.

We then performed CI to P3 heart of FOXP3^+^ Treg-infused or control NOD/SCID mice; and examined heart regeneration at day 28 post CI (Fig 4A). Our Masson’s trichrome staining showed that adoptive transfer of Treg significantly reduced cardiac fibrosis (Fig 4D, E). To investigate if heart regeneration was mediated via proliferation of cardiomyocytes, we performed immunostaining for Ki67 and cTnT (Fig 4F) and found that adoptive transfer of Treg significantly increased Ki67^+^cTnT^+^ cardiomyocytes in the border zone (Fig 4G). To evaluate heart regeneration functionally, we performed echocardiography at day 28 post CI (Fig 4H). Our data revealed that adoptive transfer of Treg significantly increased %fractional shortening compared to control (Fig 4I). We also included an additional injury model to recapitulate the role of Treg in neonatal heart regeneration. AR was performed on P3 heart of FOXP3^+^ Treg-infused or control NOD/SCID mice; and heart regeneration was examined on day 28 post injury (Fig S3A). Similarly, we demonstrated that adoptive transfer of Treg reduced cardiac fibrosis (Fig S3B) and significantly increased Ki67^+^cTnT^+^ cardiomyocytes (Fig S3C) after injury compared to control. Altogether, our gain-of-function experiments in both CI and AR models uncovered an unappreciated role of Treg in potentiating neonatal heart regeneration.

### Single cell transcriptomic profiling reveals *FOXP3*^+^ Treg as a source of paracrine factors

In order to understand how Treg regulate neonatal heart regeneration, we performed genome-wide single-cell transcriptomic profiling (scRNA-seq) as recently described^21,22^ with CD3^+^ T-cells purified at day 7 following injury of ICR mice that underwent CI at P3. In this study, the spleen served as a source of naïve T-cells (control) while the heart served as a source of neoantigen-activated T-cells. We purified about ~1850 and ~581 CD3^+^ T-cells from the spleen and heart, respectively, by flow cytometry. We then focused our analysis on *FOXP3*^+^ Treg and performed unsupervised analysis that did not rely on any known marker. *t*-distributed stochastic neighbor embedding (*t*-SNE) plots showed that Treg of spleen and heart formed distinct populations (Fig 5A): splenic Treg mainly constituted cluster C1; while heart Treg primarily formed cluster C2. We identified the most significantly upregulated genes in C1 and C2 (Table 1) and performed gene ontology (GO) functional annotations as demonstrated by pathway analyses (Fig 5B, Table S1, S2) and heatmap (Fig 5C).

**Fig. 5.**
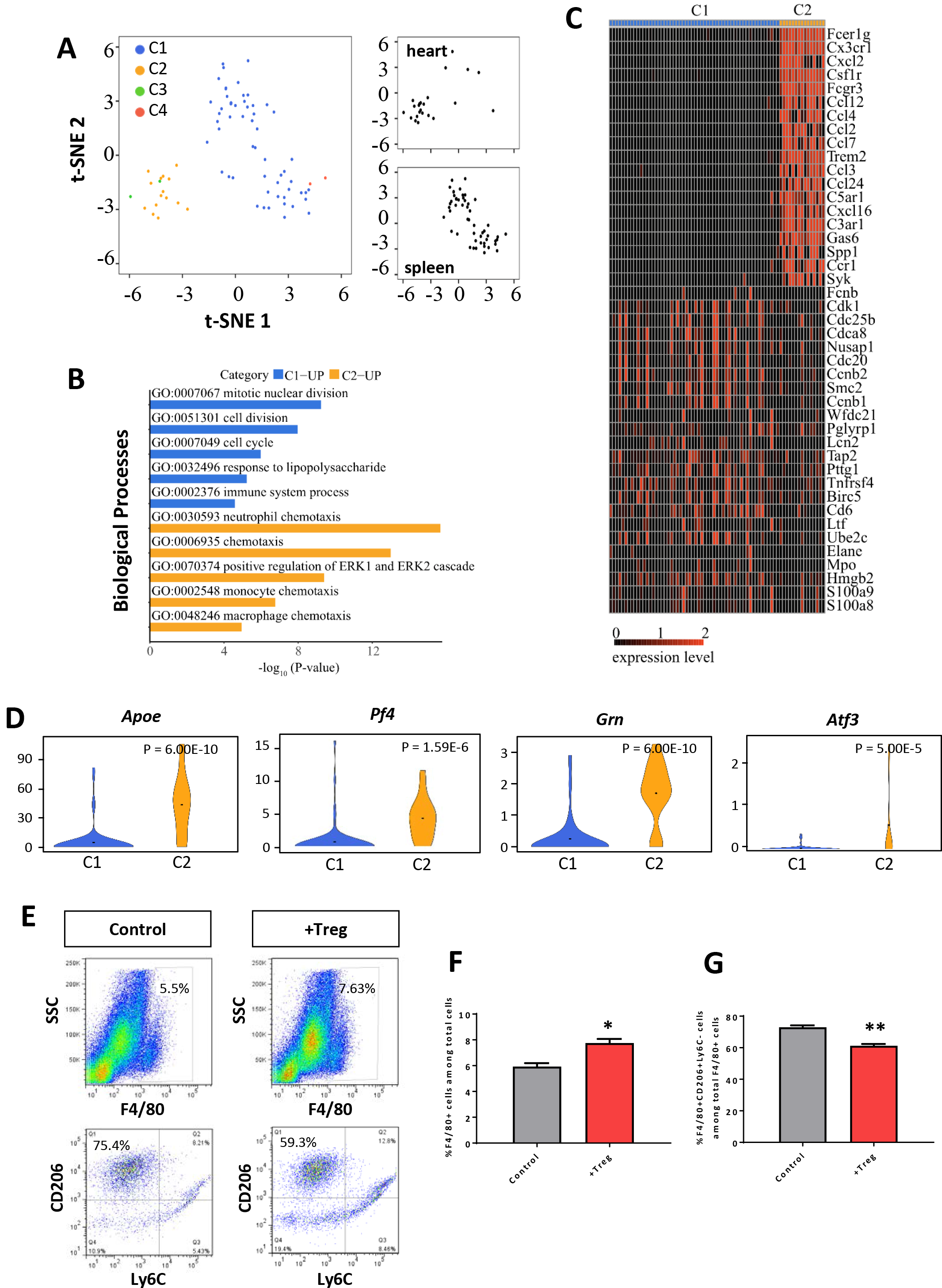
Single cell transcriptomic profiling reveals that Treg are chemoattractive to macrophages during neonatal heart regeneration. (A) Biaxial scatter plots by *t*-SNE analysis showing single-cell transcriptomic clustering of *FoxpS*^+^ Treg purified from the spleen or heart at day 7 post CI to P3 ICR mice. Cells were subgrouped into specific clusters (C1-4). (B) Analysis showing selected most significantly upregulated pathways determined by GO functional annotations in terms of biological processes of C1 splenic and C2 heart Treg (Tables S1). (C) Upregulated genes in (B) were displayed by a heatmap. (D) Violin plots showing selected most significantly upregulated genes that regulate macrophage activity or regeneration processes. (E) Flow cytometric analysis and (F, G) quantification showing (F) %F4/80^+^ macrophages among total heart cells or (G) %F4/80^+^CD206^+^Ly6C^−^ M2 macrophages among total F4/80+ macrophages at day 14 post CI with and without adoptive transfer of hCD2+ Treg to P3 NOD/SCID mice. (F, G) Data are presented as mean±S.E.M., *P<0.05, **P<0.01, n=4 per group.

**Table 1.**
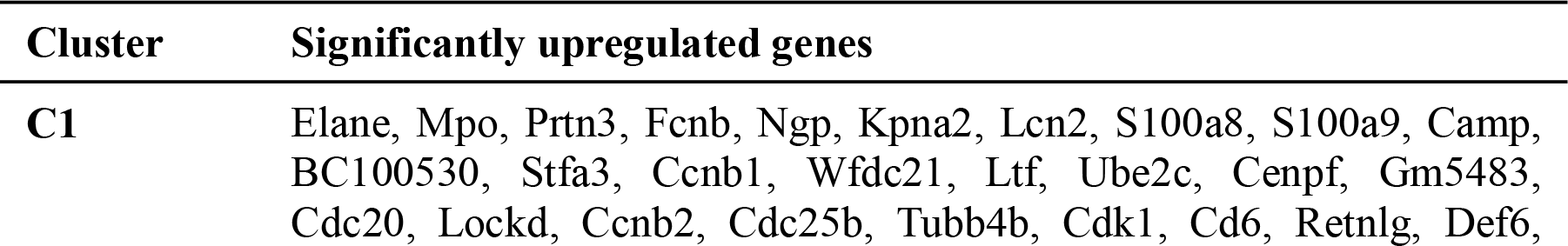
The most significantly upregulated genes expressed by heart Treg during neonatal heart regeneration. *Foxp3*^+^ Treg are purified from the spleen or heart at day 7 post CI to P3 ICR mice. C1: upregulated genes in splenic naive Treg; C2: upregulated genes in Treg following activation by neoantigens released during cryoinfarction in the heart.

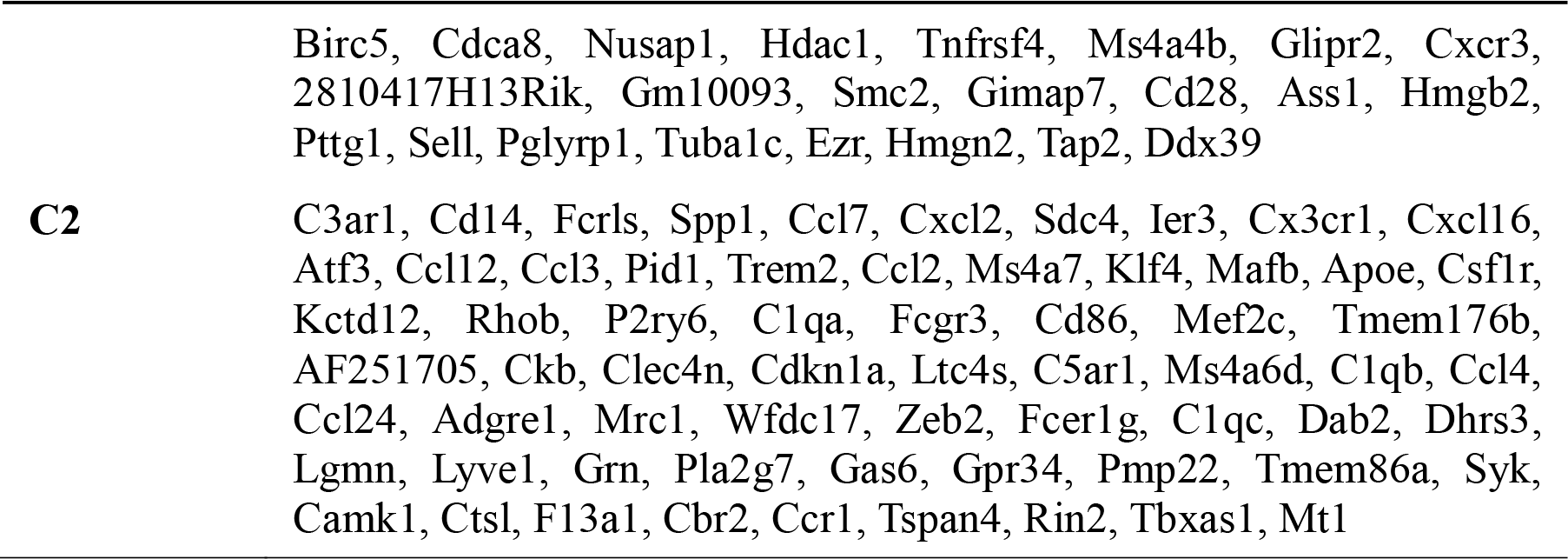

Our results revealed that C1 was distinguished by upregulated cell cycle/cell division genes, indicating that Treg of a regenerating neonatal heart were less proliferative than naive T-cells. Moreover, C2 was marked by upregulated chemotactic factors including chemokines and cytokines that attract neutrophils, monocytes and macrophages, suggesting that Treg of a regenerating neonatal heart could recruit innate immune cells including macrophages, previously reported to drive neonatal heart regeneration^4,7^. We also highlighted several significantly upregulated genes in C2 compared to C1 that have been reported to regulate macrophage activity such as apolipoprotein E (*Apoe*)^23^ and CXCL4 (*Pf4*)^25–27^; and to facilitate tissue regeneration such as granulin *(Grn)* ‘ and activating transcription factor 3 (*Atf3*)^28,29^ (Fig 5D).

To validate our scRNAseq data, we asked if Treg recruited macrophages during neonatal heart regeneration. We performed flow cytometric analysis to determine macrophage coverage in the neonatal heart of NOD/SCID mice 14 days after CI at P3 (Fig 5E). Indeed, adoptive transfer of Treg significantly increased %total F4/80^+^ macrophages (Fig 5F) yet reduced %F4/80^+^CD206^+^Ly6C^−^ M2 macrophages (Fig 5G) compared to the untreated control group. Our unbiased transcriptomic classification of FOXP3^+^ Treg demonstrated that they could be a source of paracrine factors possibly regulating neonatal heart regeneration.

### FOXP3^+^ Treg facilitate proliferation of both mouse and human cardiomyocytes in a paracrine manner

Lastly, we asked if Treg played a direct role in heart regeneration; and if they modulated heart regeneration in a paracrine manner. Since a previous report documents that Treg secreted amphiregulin (AREG) for potentiating proliferation of satellite cells during skeletal muscle regeneration^9^; and our scRNA-seq data showed significantly upregulated expression of *Atf3* in Treg during neonatal heart regeneration that has been reported to regulate *Areg* expression^30^, we hypothesized that Treg could directly facilitate heart regeneration via *Areg*. To test this, we cocultured mouse neonatal cardiomyocytes with purified hCD2^+^ Treg, supernatant of Treg cultures or recombinant mouse AREG protein for 1 day. We then performed immunostaining for Ki67 and cTnT (Fig 6A). Our results showed that hCD2^+^ Treg, supernatant of Treg or AREG significantly increased %Ki67^+^cTnT^+^ cells among total cTnT^+^ cells (Fig 6B). To recapitulate the same effect on human cardiomyocytes, we differentiated beating cardiomyocytes from human embryonic stem cells as previously described^31^ (hESC-CM, Fig 6C). We cocultured relatively mature, less proliferative hESC-CM (day 76) with supernatant of Treg cultures or recombinant human AREG protein for 3 days. Immunostaining results of Ki67 and cTnT (Fig 6D) revealed that supernatant of Treg or AREG significantly increased %Ki67^+^cTnT^+^ cells among total cTnT^+^ cells (Fig 6E). Altogether, our results uncovered that Treg directly promoted proliferation of mouse and human cardiomyocytes in a paracrine manner.

**Fig. 6.**
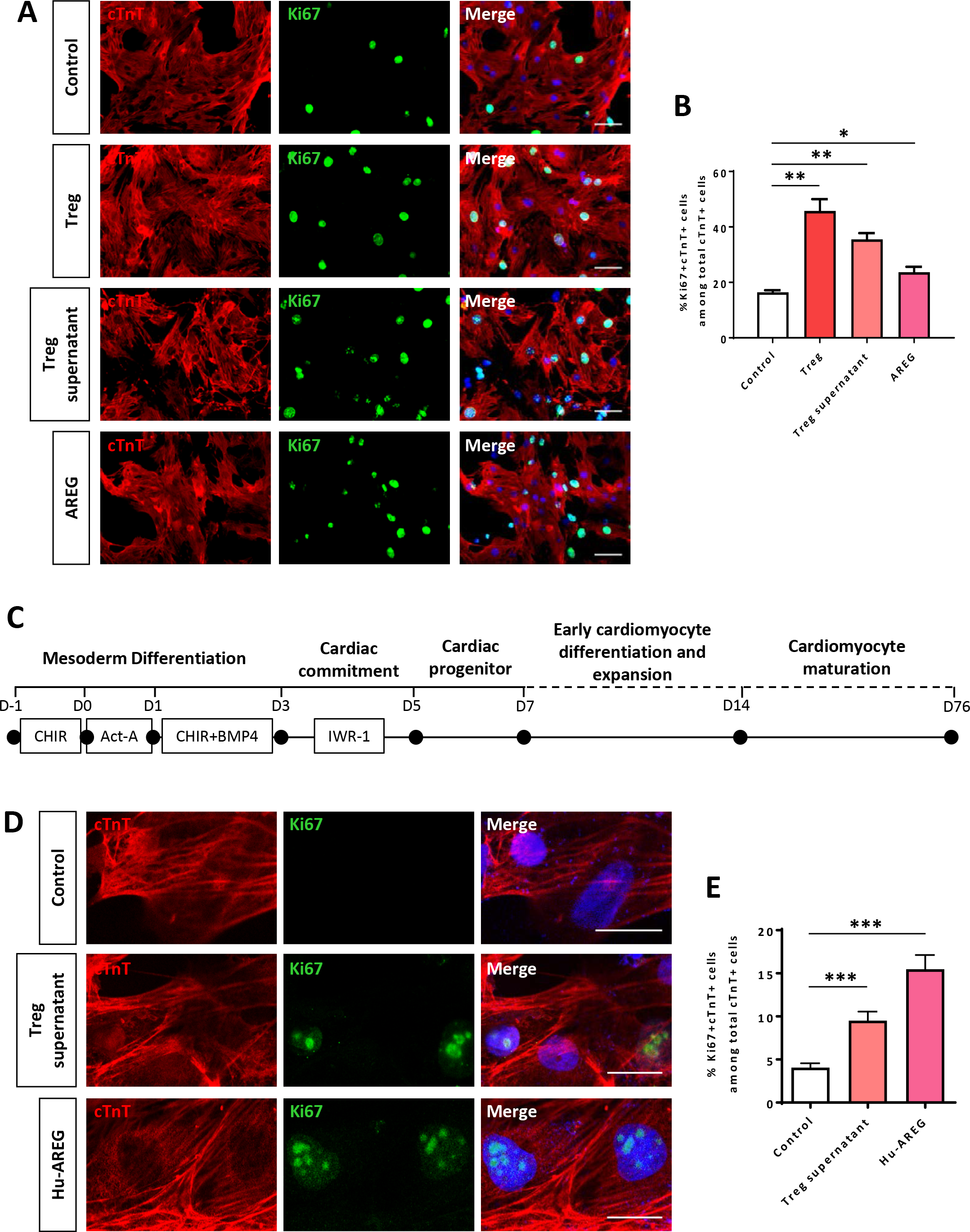
Treg directly promote proliferation of both mouse and human cardiomyocytes in a paracrine manner. (A) Immunocytochemistry for cTnT^+^ (red) and Ki67^+^ (green) cells at day 1 after coculture of CD3^+^CD4^+^hCD2^+^ Treg, Treg supernatant or mouse amphiregulin (AREG) with mouse neonatal cardiomyocytes of P1 ICR heart. (B) Quantification of %Ki67^+^cTnT^+^ proliferating cardiomyocytes among total cTnT^+^ cardiomyocytes based on (A). (C) Schematic diagram showing the differentiation protocol for generating human cardiomyocytes from embryonic stem cells (hESC-CM). (D) Immunocytochemistry for cTnT^+^ (red) and Ki67^+^ (green) cells at day 3 after coculture of Treg supernatant or human amphiregulin (AREG) with hESC-CM. (B, E) Data are presented as mean±S.E.M., *P<0.05, **P<0.01, ***P<0.001, n=3 per group.

## Discussion

Recently, immune cell-mediated tissue regeneration has been an emerging paradigm in Regenerative Biology. Macrophages, in particular, are the most studied immune cell type for tissue repair and regeneration. Depletion of macrophages in mice contributes to excessive cardiac fibrosis, impeded neoangiogenesis and consequently impaired neonatal heart regeneration after injury^4,7^. In fact, adaptive immune cells including T-cells often regulate macrophage activation during immune responses; whether T-cells influence neonatal heart regeneration is, however, not investigated.

In this study, we examined if adaptive immune cells particularly T-cells mediate neonatal heart regeneration after injury because our results showed that innate immune cells including macrophages were insufficient to drive functional regeneration without the presence of T-cells in NOD/SCID mice. We then asked which subset of T-cells is important for neonatal heart regeneration. In wile type mice, we found that there was no significant infiltration of CD8^+^ T-cells into the infarcted myocardium after CI compared to that of the sham-operated control; and depletion of which did not alter the healing processes, suggesting that CD8^+^ T-cells were not likely required for neonatal heart regeneration. Instead, we found significant infiltration of CD4^+^ T-cells after CI to a P8 heart that was unable to regenerate compared to the sham-operated control. Depletion of CD4^+^ T-cells via the lytic GK1.5 antibody leads to significantly reduced cardiac fibrosis and enhanced cardiomyocyte proliferation of a P8 heart after CI, indicating that CD4^+^ T-cells negatively regulate neonatal heart regeneration. Unexpectedly, we found a population of CD4^−^FOXP3^+^ cells following GK1.5 treatment. Unlike conventional CD4^+^ T-cells, FOXP3^+^ Treg can recognize antigens without CD4 as they are able to utilize other mechanisms such as LAG3 that further enhances their suppressor function^32^. Therefore, GK1.5 treatment could inhibit function of conventional CD4^+^ T-cells yet preserve function of CD4^−^FOXP3^+^ Treg, indicating that FOXP3^+^ Treg might play a role in heart regeneration.

Recently, accumulating evidence shows that FOXP3^+^ Treg exert a direct or indirect role in tissue repair and regeneration. For instances, Burzyn et al. reveal that Treg directly promote activation of satellite cells and regeneration of skeletal muscle via secretion of amphiregulin^9^; Arpaia et al. demonstrate that Treg facilitate tissue repair after influenza infection^12^; Dombrowski et al. unveil that Treg promote differentiation of oligodendrocytes and regeneration of myelin in the central nervous system^14^; and Ali et al. uncover that Treg facilitate proliferation and differentiation of hair follicular stem cells in skin^11^. More recently, we have also showed that Treg promote regeneration of blood vessels in the peripheral artery system after ischemic injury in type-2 diabetes^15^. In fact, targeting T-cells for tissue regeneration offers unappreciated advantages such as specificity towards autoantigens released during injury. Indeed, we have observed that the therapeutic effect of Treg in vascular regeneration was only localized in the ischemic but not non-ischemic tissue^15^.

To ask if Treg facilitate neonatal heart regeneration, we performed both loss- and gain-of-function studies. Depletion of CD4^+^CD25^hi^FOXP3^+^ Treg via the lytic anti-CD25 antibody after CI to a regenerating P3 heart of NOD mice led to impaired heart regeneration manifested by significantly increased cardiac fibrosis and reduced number of proliferating cardiomyocytes compared to the untreated control. For gain-of-function experiments, we generated the *Foxp3*^hCD2^ reporter mice in NOD background as a source of hCD2^+^ Treg for adoptive transfer into NOD/SCID mice. Engraftment of Treg contributed to significantly reduced cardiac fibrosis, increased number of proliferating cardiomyocytes and enhanced cardiac function 4 weeks after CI to a non-regenerating P3 heart of NOD/SCID mice compared to control. Similarly, adoptive transfer of Treg also promoted neonatal heart regeneration of NOD/SCID mice in an additional injury model, AR.

Mechanistically, single cell transcriptomic profiling revealed that Treg of a regenerating P3 heart were a source of chemokines and cytokines that could recruit monocytes and macrophages, previously reported to drive neonatal heart regeneration. We validated this by flow cytometric analysis on macrophages 2 weeks after CI to a P3 heart of NOD/SCID in the Treg-infused group compared to that of the Treg-deficient control. Indeed, our results showed that adoptive transfer of Treg significantly increased %F4/80^+^ macrophages in the infarcted myocardium of NOD/SCID mice. Nevertheless, there was a significant reduction in %F4/80^+^CD206^+^Ly6C^−^ M2 macrophages after adoptive transfer of Treg. In fact, M2 macrophages have been reported to be correlated with cardiac fibrosis via secretion of TGFβ, osteopontin and galatin-3^33,34^

In addition to orchestrating macrophages, our scRNAseq experiments also demonstrated that Treg of a regenerating P3 heart significantly enhanced expression of genes found in regeneration of other organ systems such as *Grn*^25–27^ and *Atf3*^28,29^ compared to naïve Treg. Intriguingly, *Atf3* has been documented to regulate expression of *Areg*^30^ previously shown to promote Treg-mediated satellite cell proliferation and skeletal muscle regeneration^9^. Since proliferation of cardiomyocytes is the major mechanism by which a neonatal heart regenerates, we asked if Treg or Treg secreted factors such as AREG has a direct role in proliferation of cardiomyocytes. Our coculture experiments demonstrated that supernatant of Treg or AREG promoted both mouse and human proliferation of cardiomyocytes, indicating that Treg could regulate heart regeneration in a paracrine manner. Taken together, we have highlighted a hitherto novel role of Treg in potentiating neonatal heart regeneration by regulating macrophage activity and facilitating cardiomyocyte proliferation after injury. Our findings resolve a long-standing question about how the neonatal heart regenerates; and offer new avenues for developing novel therapeutics targeting Treg-mediated heart regeneration.

## Materials and Methods

### Mice

*Foxp3*^hCD2^ reporter mice (C57BL/6), in which expression of the surface hCD2 molecule is driven under the *Foxp3* promoter^18^, were backcrossed onto the NOD/ShiLtJ (Clea Japan. Inc) background for 12 generations. Males were sacrificed for analysis at 4-8 weeks old before onset of diabetes. All animal procedures were approved by the CUHK Animal Experimentation Ethics Committee and performed in compliance with the *Guide for the Care and Use of Laboratory Animals* (NIH publication, eighth edition, updated 2011).

### Neonatal mouse heart cryoinfarction (CI)

CI was performed as previously described^17^. Briefly, neonatal mice at P3 or P8 were subjected to anesthesia by freezing for ◻3-5 minutes, and were then placed on the frozen operation table once breathing was steady. Mouse limbs were fixed in location with forceps and 70% ethanol was applied to disinfect the surgical area. An incision (◻1 cm) was made along the sternum and vertical of the chest muscles under a stereomicroscope. Two or three intercostal incisions were made in the left sternum chest to separate the pericardium and expose the left ventricle. Blunt port copper wire (1 mm thickness) was frozen in liquid nitrogen and then put on the left ventricle in order to induce frostbite; this was maintained for ◻7-8 seconds until the left ventricle appeared white. After injury, bubbles and blood in the chest were squeezed out. The chest was closed and the skin was sewn up with 8-0 sutures. After surgery, neonatal mice were placed under a 37°C heating pad to keep warm. They were then placed back with their mothers as soon as they woke up and the skin color returned to normal. In the sham-operated control, we performed the same experimental procedures as above except that we replaced liquid nitrogen with PBS at room temperature.

### Neonatal mouse heart apical resection (AR)

AR was performed as previously described^17^. Briefly, neonatal mice at P3 were subjected to anesthesia by freezing for 3-5 minutes, and were then placed on the frozen operation table once breathing was steady. Mouse limbs were fixed in location with forceps and 70% ethanol was applied to disinfect the surgical area. An incision (◻1 cm) was made along the sternum and vertical of the chest muscles under a stereomicroscope. Two or three intercostal incisions were made in the left sternum chest to separate the pericardium and expose the apex. Curved forceps were extended into the intrathoracic to pull out the heart and the apex was then truncated with microsurgical scissors. After injury, bubbles and blood in the chest were squeezed out. The chest was closed and the skin was sewn up with 8-0 sutures. After surgery, neonatal mice were placed under a 37°C heating pad to keep warm. They were then placed back with their mothers as soon as they woke up and the skin color returned to normal. In the sham-operated control, we performed the same experimental procedures as above except that we did not truncate the heart apex.

### Adoptive transfer of Treg

Treg were purified from the spleen of 4-6 weeks old *Foxp3*^hCD2^ mice using anti-hCD2 magnetic beads following the manufacturer’s instructions (Miltenyi Biotech). Each neonatal NOD/SCID mouse was adoptively transferred with 1 million Treg through intraperitoneal (i.p.) injection immediately after CI, AR or sham surgery.

### Administration of mAb

Lytic anti-CD4 (clone GK1.5, BioxCell), lytic anti-CD8 (clone YTS169, BioxCell) and lytic anti-CD25 (clone PC61, BioxCell) mAbs were used as previously described^19,35^. Briefly, YTS177 was injected i.p. at 0.4 mg/pup on days 0, 7, 14 and 21 following CI; and GK1.5 or YTS169 was injected i.p. at 0.2 mg/pup on days 0, 2, 4, 6 and 10 following CI. PC61 was injected i.p. at 0.5 mg/pup on days 0, 2, 4, 6 and 10 following CI.

### Echocardiography

The left ventricle systolic function was measured 2 months following CI or sham operation with echocardiography in mice via a digital ultrasound system (Vevo2100 Imaging System, VisualSonics). Conventional measurements of the left ventricle (LV) included: end-diastolic diameter (LVEDD), end-systolic diameter (LVESD), intraventricular septal thickness (IVST), posterior wall thickness (LVPWT), ejection fraction (LVEF) and fractional shortening (LVFS). Analysis was performed with VisualSonics Vevo2100 software.

### Immunostaining

Heart tissues were dissected and fixed in 4% paraformaldehyde at 4°C overnight. The fixed tissues were washed three times with PBS and equilibrated in 30% sucrose at 4°C for 2 days before freezing and cryosectioning. Eight micrometer frozen sections were blocked at 2% goat serum and then stained with the respective primary antibodies at 10 ug/ml at 4°C overnight. Anti-mouse primary antibodies used: COLA1 (abcam, ab34710), cTNT (abcam, ab8295) and Ki67 (eBiosciences 14-5698-82). Anti-human primary antibodies used: cTNT (RnD systems, MAB1874) and Ki67 (abcam, ab 15580). Alexa-Fluor-488− or Alexa-Fluor-546-conjugated secondary antibodies (Invitrogen) were used at room temperature for 30 minutes in the dark. Slides were mounted with DAPI-containing fluorescence mounting medium (Dako) and fluorescence was detected with an upright fluorescence microscope, inverted fluorescence microscope or confocal microscope (all Leica). Images were processed with ImageJ software and cTNT coverage was analyzed based on this formula: cTNT^+^ area/total area.

### Masson’s trichrome staining

Masson’s trichrome staining was performed to determine collagen deposition per manusfacturer’s instruction (Polysciences 25088-1). Briefly, frozen sections were washed in PBS and fixed in 4% paraformaldehyde for 8 min. Slides were then incubated in Bouins’ solution (5% acetic acid, 9% formaldehyde and 0.9% picric acid) at room temperature overnight. The next day after washed with distilled water, slides were incubated in Weigert’s iron hematoxylin solution for 10 minutes, washed and then stained with Biebrich scarlet-acid fuchsin for 5 minutes. After three washes with distilled water, slides were incubated in phosphotungstic/phosphomolybdic acid for 10 minutes followed by staining with aniline blue solution for 5 minutes. After that, slides were washed with distilled water for three times and dehydrated with ethanol and xylene based on standard procedures. Images were acquired on Nikon eclipse TE2000-S microscope. For the CI group, images were analyzed and fibrosis coverage was quantified based on this formula: scar perimeter/total perimeter.

### Flow cytometry, cell sorting and analysis

To collect splenocytes, the excised spleen was dissociated in PBS with a syringe plunger through a 40um cell strainer to obtain single cell suspension. To study immune cell infiltrates in the neonatal heart, heart tissues were minced into small fragments and dissociated with 1:1 type II collagenase (1000 U/ml in PBS, Worthington) and dispase (11 U/ml in PBS, Gibco) at 37°C for 30 minutes. Enzymatic action was stopped by adding 10% FBS and the dissociated cells were washed twice with PBS. The dissociated single splenocytes or neonatal heart cells were removed from the contaminated erythrocytes by incubating with the red blood cell lysis buffer (eBiosciences) for 5 minutes; and were then blocked with 2% normal rabbit serum. Cells were subsequently stained with fluorochrome-conjugated antibodies against the following antigens: mCD3, mCD4, mCD8, mCD31, mCD45, mF4/80, mLy6C or hCD2 (Biolegend or eBiosciences) at a dilution of 1:100, unless specified by the manufacturer, at 4°C for 30 minutes. Murine Treg were detected with the Treg staining kit according to manusfacturer’s instructions (eBioscience). Cells were then washed three times with 2% FBS-containing PBS and analyzed on flow cytometer (BD FACSAria™ Fusion). Propidium iodide (PI, BD) positive dead cells were excluded for live cell analysis/sorting; and FACS data were then analyzed with the FlowJo software (Tree star).

### Cell cultures

For Treg, näive CD3^+^CD4^+^hCD2^+^ Treg were purified from the spleen of 4-6 week old *Foxp3*^hCD2^ reporter mouse by flow cytometry and were stimulated before cocultured with cardiomyocytes as described previously^36^. Briefly, each well of a 96-well plate was coated with 10 ug/ml anti-CD3 (Biolegend, 100314) and 1 ug/ml anti-CD28 (Biolegend, 102112) at 4°C overnight. The plate was then washed with PBS twice. Treg were cultured *in vitro* with RPMI1640 supplemented with 10% heat inactivated fetal bovine serum, 1% sodium pyruvate (Life Technologies), 10 mM HEPES (Life Technologies), 50 uM 2-mercaptoethanol (Life Technologies), 40 ng/ml IL-2 (Peprotech, 212-12) and 10 ng/ml TGFb (RnD systems, 7666-MB-005) at 37°C for 4 days before coculture experiments.

For murine neonatal cardiomyocytes, they were isolated with an enzymatic digestion approach as previously described^37^. Briefly, P1 ventricles were minced into small fragments and pre-digested in 0.05% trypsin-EDTA at 4°C overnight. The pre-digested mixture was washed with 10 mM HEPES and 1X penicillin/streptomycin-containing DMEM/F12 medium (light medium) pre-warmed at 37°C, followed by repeated digestions in a stepwise manner: the tissues were digested with 100 U/ml type II collagenase at 37°C for 10 minutes. After that, the supernatant was collected and mixed in a ratio of 1:1 with DMEM/F12 medium supplemented with 10mM HEPES, 1X penicillin/streptomycin, 10% horse serum (Invitrogen) and 5% fetal bovine serum (dark medium). The supernatant mixture was then kept on ice and the tissue pellet was further digested for 2-3 times with the same procedures until the tissues became single cells. All supernatant mixtures were then pooled together and centrifuged at 800 rpm for 5 minutes. Differential plating was performed to remove fibroblasts by resuspending the cell pellet with 10 ml dark medium followed by seeding onto a T25 flask at 37°C for 1 hour. After that, the unattached cells were transferred to a new T25 and replated at 37°C for another hour. The unattached cardiomyocytes were then centrifuged at 400 rpm for 5 minutes, and resuspended with appropriate volume of dark medium for cell counting. Cardiomyocytes were plated on Matrigel (1:100 in DMEM/F12)-coated chamber slide at a density of 10,000 cells per well and cultured in dark medium at 37°C for 24 hours. To synchronize proliferation of cardiomyocytes before experiment, they were starved with serum-free medium overnight. After that, they were cocultured with *in vitro* stimulated Treg in a ratio of cardiomyocytes: Treg as 3:1, Treg supernatant-containing dark medium (1:1) or 50 ng/ml murine recombinant amphiregulin (RnD systems, 989-AR-100) at 37°C for 1 day before analysis.

For human cardiomyocytes, cardiomyocytes were derived from hESCs as previously described^31^. Cardiomyocytes were generated with a sequential administration of growth factors (Fig 6C). Briefly, hESCs were seeded at a density of 3.75 × 10^4^ cells/cm^2^ on matrigel (BD, growth factor reduced, 1:60)-coated plate in mTeSR™ (Stemcell Technologes) supplemented with 10 uM Y-27632 (Selleck Chem, S1049) and 1uM CHIR99021 (Selleck Chem,S2924). On the day of mesoderm induction, cells were overlaid with Matrigel (1:60) and 100 ng/ml Activin A (Peprotech, 120-14E) in insulin-free B27-containing RPMI1640 medium (RPMI/B27-) for 24 hours, followed by 1uM CHIR99021 and 5 ng/ml BMP-4 (Peprotech, AF-120-05ET) for 48 hours. Subsequently, the medium was refreshed and supplemented with 5 uM IWR-1 (Calbiochem, 681669) for 48 hours, followed by unsupplemented RPMI/B27-medium for 48 hours. On day 7, cardiac progenitors were expanded and maintained in insulin^+^ B27-containing RPMI1640 medium (RPMI/B27+). The medium was refreshed every 3 days. Beating cardiomyocytes could often be observed on day 14 of differentiation. To synchronize proliferation of cardiomyocytes, beating cardiomyocytes were starved in RPMI1640 alone for 9 hours followed by replenishment with RPMI/B27+ alone, supernatant of Treg cultures or 50 ng/ml human recombinant amphiregulin (Peprotech, 100-55B) for 3 days before analysis.

### Single-cell encapsulation and library preparation

Single cells were purified by FACS sorting before library preparation and single-cell libraries were prepared with the Chromium Single Cell 3’ Reagent Kits v2 (10x Genomics) as per manufacturer’s instructions. Briefly, sorted cells in suspension were first prepared as gel beads in emulsion (GEMs) on Single Cell 3’ Chips v2 (10x Chromium) using the Chromium Controller (10x Genomics). Barcoded RNA transcripts in each single cell were reverse transcribed within GEM droplets. cDNA was purified with DynaBeads MyOne Silane beads (Invitrogen) and then amplified for subsequent library construction. Sequencing libraries were prepared by fragmentation, end-repair, ligation with indexed adapters and PCR amplification using the Chromium Single Cell 3’ library kit v2 (10x Genomics). Nucleic acid was cleaned up after each steps using SPRIselect beads (Beckman Coulter). Libraries were then quantified by Qubit and realtime quantitative PCR on a LightCycler 96 System (Roche).

### Single-cell RNA-sequencing and Functional Annotations

Pooled libraries were sequenced on the Illumina HiSeq 2500 platform. All single-cell libraries were sequenced with a customized paired-end dual index format (98/26/0/8 basepair) according to manufacturer’s instructions. Data were processed, aligned and quantified using the Cell Ranger Single-Cell Software Suite (v 2.1.1)^38^. Briefly, data were demultiplexed based on the 8 base-pair sample index, 16 base-pair Chromium barcodes and 10 base-pair unique molecular identifiers (UMI). After quality control, reads were aligned on *Mus Musculus* Cell Ranger transcriptome reference (mm10-1.2.0). Data analyses, including tSNE and graph-based clustering, were performed according to Cell Ranger’s pipelines with default settings. Differentially expressed genes in each cluster relative to all other clusters were identified by Cell Ranger’s pipelines with default settings (minimum mean expression = 1 and adjusted p-value cutoff = 0.05). The top N genes by log2-fold change for each cluster were further analyzed (N=10000/K^2, where K is the number of clusters). Gene ontology (GO) enrichment analyses of the cluster-specific highly expressed genes were performed by DAVID Bioinformatics Resources (v6.8)^39^.

### Statistical analysis

The data were expressed as arithmetic mean±S.E.M. of biological replicates (n = 5, unless otherwise specified) performed under the same conditions. Statistical analysis was performed using the unpaired student’s t-test with data from two groups; while date from more than two groups was performed using an ANOVA followed by Tukey’s method for multiple comparisons. Significance was accepted when *P* < 0.05.

## Acknowledgments

We thank Dr. Joaquim S.L. Vong and Xisheng Li (CUHK) for their technical help during this study. This work was supported by Research Grants Council of Hong Kong (24110515, 14111916, C4024-16W, C4026-17WF); Health and Medical Research Fund (03140346, 04152566); Croucher Foundation (Innovation Award and Start-up Allowance); Direct Grant, Faculty Innovation Award, Seed Fund from Lui Chi Woo Institute of Innovative Medicine, postdoctoral fellowships (K.Y.Y. and R.C.Y.T) and postgraduate studentship (J.L.) from CUHK.

## Author contributions

J.L., R.C.Y.T., V.C. and B.S. performed experiments; B.Z. and Y.M.D.L. contributed reagents; J.L., K.Y.Y., R.C.Y.T. and K.O.L. analyzed data; K.O.L. designed research and wrote the manuscript.

